# High-Precision Event Synchronization for Chronic Deep Brain Stimulation Local Field Potential Recordings

**DOI:** 10.64898/2026.05.13.724854

**Authors:** Sophia Gimple, Yasin Temel, Christian Herff, Marcus L.F. Janssen

## Abstract

**Background:** Electrophysiological recordings from chronically implanted Deep Brain Stimulation (DBS) electrodes can greatly advance understanding of disease and treatment mechanisms of motor and psychiatric disorders. The Medtronic Percept system allows for chronic recordings of local field potentials (LFP) from DBS target regions. However, these systems lack an inbuilt synchronization option to align LFP recordings to other recording modalities and consequently events in computerized tasks.

**Objective:** We propose and evaluate a synchronization method based on Transcutaneous Electrical Stimulation (TES) with low amplitudes to precisely align recorded LFP signals from the DBS electrodes to EEG recordings.

**Methods:** The TES-based synchronization approach was implemented and tested in 11 participants implanted with the Medtronic Percept for treatment of Parkinson’s disease.

**Results:** The proposed method provides high reliability, precise alignment and usability across all Medtronic Percept recording modes. Notably, the method enables recordings during adaptive DBS and with stimulation turned off. In this recording mode, LFP signals can be acquired from all recording contact pairs simultaneously, with a high signal-to-noise ratio. We provide detailed setup plans and share Python and Matlab scripts for signal alignment to enable easy application of our approach.

**Conclusion:** By enabling reliable, well-aligned LFP recordings from all DBS contacts, our method provides a robust tool for studying neural dynamics and refining therapeutic interventions in diverse neurological conditions.

## 1. Introduction

Deep brain stimulation (DBS) of the basal ganglia is a well-established treatment for motor symptoms in movement disorders such as Parkinson’s disease [1]. DBS is also increasingly applied in treatment-resistant psychiatric disorders such as OCD, depression and Tourette syndrome [2, 3, 4]. Chronic deep brain stimulation involves implanting electrodes in the basal ganglia connected to an implantable pulse generator (IPG) powering and steering the stimulation. Several DBS systems are currently both CE- and FDA-approved and used clinically. Among these systems, the Medtronic Percept(TM) and Newronika AlphaDBS(TM) system stand out by enabling neurophysiological recordings of local field potential (LFP) measurements from DBS electrode contacts [5, 6]. LFP recordings offer the unique opportunity to study basal ganglia processes and their relation to disease symptoms and treatment mechanisms [5]. However, aligning LFP signals to behavioral events or other measurements remains challenging [7, 8]. Precise alignment is essential to capture the temporal dynamics of neural signals time-locked to events, supporting reliable interpretation and biomarker discovery from LFP recordings [9]. Previous papers have repeatedly described the need for external synchronization strategies that can compensate for the lack of an in-built synchronization option in the system and proposed a variety of possible solutions (e.g. [7, 10, 11, 12]). Here, we propose and evaluate a standardized synchronization option based on low-amplitude Transcutaneous Electrical Stimulation (TES) applicable across all Percept Medtronic recording modes.

### 1.1. Recording modes of the Medtronic Percept

The Medtronic Percept enables continuous LFP recordings at 250 Hz in two different recording modes, summarized in Table 1. Generally, each electrode consists of 4 contact rings (labeled 0-3 from bottom to top, see Figure 3).In *BrainSenseTimeDomain* Streaming mode, LFP signals are recorded from one electrode ring pair per hemisphere, which must, due to Software constraints, be positioned directly above and below the stimulation contact(s). This allows for recordings from contact pairs 0-2 (stimulation at contact ring 1), 0-3 (stimulation at 1 and 2) or 1-3 (stimulation at 2). While the legacy leads have 4 uninterrupted rings with one contact each, in the Sensight directional leads rings 1 and 2 are subdivided into three contacts (1a,1b,1c and 2a,2b,2c respectively). Using the On/Off button, stimulation can also be switched off. Stimulation On at 0mA and Stimulation Off (by definition at 0mA) do however not result in equivalent LFP recordings. Stimulation On *BrainSenseTimeDomain* recordings may exhibit ECG artifacts and stimulation artifacts not visible in Stimulation Off recordings [13, 14, 15]. The second recording mode, *IndefiniteStreaming*, allows for recording LFP signals from 3 contact pairs per electrode simultaneously (0-2, 0-3 and 1-2). Stimulation must be switched off, resulting in the reduction of ECG artifacts, as also observed in *BrainSenseTimeDomain* with Stimulation Off.

**Table 1.** Overview of Recording Modes and Technical Constraints

| Recording mode | Number of recording contact pairs | Possible ECG artifacts | Stimulation artifacts | DBS stimulation possible | Conditions for recordings |
| --- | --- | --- | --- | --- | --- |
| BrainSenseTimeDomain (ON)* | 1 per lead | Yes** | Yes | Yes | Stimulation at contacts 1, 2, or 1+2. |
| BrainSenseTimeDomain (OFF) | 1 per lead | No | No | No | Stimulation turned off. |
| IndefiniteStreaming | 3 per lead | No | No | No | Stimulation turned off. |

**Table 2.** Medtronic Percept recording modes. In *BrainSenseTimeDomain* mode recordings can be performed either with stimulation turned On (0mA possible) or stimulation turned off. In *IndefiniteStreaming* mode recordings can be performed from 3 recording contact pairs under stimulation off. *please note that stimulation amplitude can be set at 0mA; **ECG artifacts may be present

### 1.2. Synchronization approaches

Previous research has described a number of possible synchronization methods. Generally, in these methods an artifact is induced that can be detected both in the LFP recordings and by another recording modality such as a simultaneous EEG, MEG or video recording. Commonly, EEG or MEG recordings comprise of an inbuilt synchronization method e.g. real-time markers that can be triggered by a computerized task. Synchronizing the LFP data to the EEG or EMG data therefore allows for precise alignment of the LFP data to task-events. Previously described synchronization methods utilize artifacts induced by rapidly switching DBS stimulation amplitude [7, 16, 17, 10], tapping on the IPG [18, 12, 17], and alignment based on ECG artifacts [19]. Furthermore, artifacts for synchronization can be induced using transcutaneous electrical stimulation (TES), commonly applied for nerve stimulation (TES). This method is especially promising for precise synchronization in stimulation off recordings and has previously been piloted by Thenaisie et al. in two test participants, reporting visible artifacts in all six recording channels with a synchronization accuracy of 12ms [7]. Furthermore, Alarie et al. describe event-linked automated TES stimulation using short stimulation bursts in two participants. They report worse synchronization outcomes under stimulation off with artifact jitter of ±22.9 ms compared to stimulation On at ± 9.08 ms [20]. However, safety and feasibility of synchronization based on TES artifacts has, to the best of our knowledge, not been formally described and evaluated in a larger study population. The aim of this study was to formally evaluate synchronization between LFP and EEG signals using TES artifacts in 10 participants and show feasibility of applying this method for alignment in the recordings of a computerized cognitive task.

## 2. Methods

### 2.1. Participants

Patients, diagnosed with PD and treated with DBS with a Percept (Medtronic Percept™ PC Neurostimulator), were included between April 2025 and January 2026. Participants were excluded if TES at 1.5mA was perceived as unpleasant. There were no further exclusion criteria. In total, 11 out of 14 evaluated patients were included (Mean age 62.2 (std=7.81, median=63), 4 female). Ten participants were implanted with the SenSight™ Directional Leads, one with Medtronic Legacy Leads. IPG location was either subclavicularly in the right chest (5) or in the right abdomen (6 participants). Written informed consent was obtained from all participants. The study was waived by the medical ethical committee of the Maastricht University Medical Centre (MUMC+) in Maastricht, The Netherlands (Protocol number 2023-0140).

### 2.2. Recording setup

LFP signals were recorded from the Medtronic percept device with a sampling rate of 250Hz. For recordings during stimulation, LFPs were measured using the *BrainSenseTimeDomain* recording mode in combination with clinical stimulation parameters. At time points, when clinical treatment required the stimulation to be turned off, additional measurements were performed in the *BrainSenseTimeDomain* mode under stimulation on with stimulation amplitude set to 0mA and under stimulation off. Additional stimulation-off recordings were conducted in the *IndefiniteStreaming* mode. EEG was acquired using a cap with 21 electrodes, placed according to the international 10–20 system, with Braintronics amplifiers and BrainRT software at 256 Hz sampling rate (OSG, Belgium, v4.03.00).

### 2.3. TES-based synchronization setup

For synchronization of LFP and EEG recordings, TES artifacts were induced using a medical Nicolet EDX system with Natus Elite - Synergy software version 22.4 (Natus neurology Inc., Middleton WI, USA). TES stimulation was delivered via two Ag/AgCl-coated cup electrodes filled with Ten20 conductive paste and positioned on both mastoids directly behind the ears beneath the EEG cap (compare Figure 1 A). The anode and cathode were assigned randomly. The skin was prepared using Nuprep skin prep gel. TES was applied at 80 Hz with a pulse width of 0.5 ms. The amplitude was titrated individually for each participant. Starting at 0mA, TES Stimulation amplitude was increased 0.25 mA steps. Amplitude was selected to be as high as possible while not being perceived as unpleasant by participants. The maximal selected stimulation intensity was 1.96 mA.

**Figure 1.**
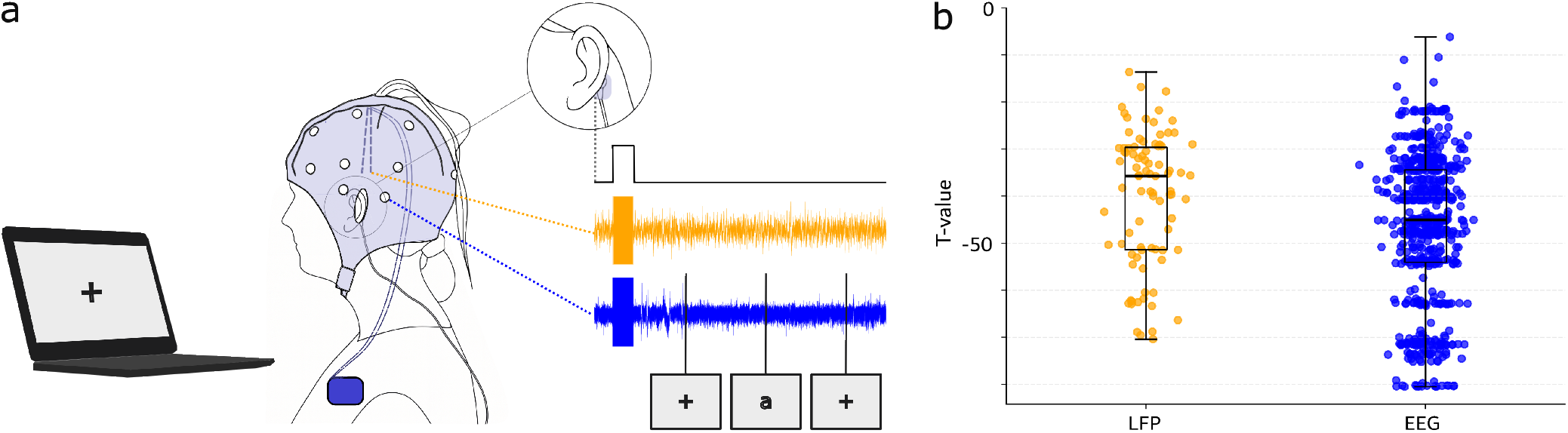
a) Recording and synchronization setup: The participant performs a computerized task while LFP (orange) and EEG signals (blue) are measured from the implanted DBS electrodes and the superficial EEG electrodes respectively. Markers representing specific task events (e.g. fixation cross or stimulus presentation) are saved along with the EEG data. On the mastoids behind the ears of the participants, additional electrodes connected to a stimulation device are placed. TES stimulation is switched on and off before and after the task to induce artifacts both in EEG and LFP recordings, which can be used for synchronization between the two electrophysiological signals. b) TES stimulation induces artifacts in a specific frequency band (in our case 80Hz). T-Values of 75-85 Hz power during intervals with and without artifacts of LFP and EEG signals. Each dot represents one electrode during one recording session.

### 2.4. LFP signal processing

When using the Medtronic Percept system, recording data is saved in packages with corresponding time-stamps at a sampling rate of 250Hz. Multiple preprocessing steps were performed to ensure high data quality. Package sizes and time-stamps were used to check for erroneous data transfer. Missing data were interpolated using local averaging within a ± 300 ms window. If packages exceeded the expected size, excess data was removed to fit the expected package size. Too big packages at the beginning of the recording were merged and trimmed from the start. If too big packages occurred later during the recording, samples from the end of the too big package were dropped. LFP signals were resampled from 250 to 256 Hz to match the EEG sampling rate. As we induced TES artifacts with a frequency of 80Hz, signals were bandpass-filtered between 75 and 85 Hz ( 3rd-order Butterworth). Afterwards, the absolute value of the Hilbert transform of the filtered signal was extracted.

### 2.5. Characterization of induced artifacts

In a first step, we evaluated the success rate of inducing significant artifacts in the Hilbert transformed signal in both EEG and LFP recording channels. We compared the time intervals including artifacts with the time intervals before and after the artifact time-window combined. Using independent t-tests, we tested for a difference in power in the 75 to 85Hz frequency band between the artifact and no-artifact time-windows. We performed Bonferroni correction to account for multiple testing. We expected artifact-segments to show a significantly higher power compared to no-artifact segments, leading to a significant t-test result with negative t-value.

### 2.6. LFP and EEG alignment procedure

To align LFP and EEG recordings, we used a cross-correlation approach selecting the channel pair with the highest overall correlation score. Filtered signals of both recording modalities were used (compare Figure 2 a, b, c). Recordings from all LFP and EEG channels were candidate signals for synchronization. In a multi-step procedure, we iterated over all LFP and EEG channel pairs. For each LFP-EEG channel pair, we computed cross-correlation scores. The channel pair with the highest overall correlation score was selected and the corresponding offset described our best estimate of time offset between the LFP and EEG signal. In a next step, signals were aligned using the estimated offset. (compare Figure 2 b and d). In case of multiple artifacts, we could summarize them into different groups and use subparts of the signal to align those independently. We used groups of at least three artifacts to allow for multiple on and off-switching flanks that were used for cross-correlation. Offsets computed from separate parts of the signal with independent artifact groups were compared to estimate the consistency of alignment (compare Figure 3a). For final alignment, we propose to use the mean of the computed offsets, but other options like selecting the offset of the artifact group with the highest correlation are also possible. To ensure transparency and reproducibility, we provide the Python and MATLAB scripts used for signal alignment (https://github.com/GimpleSophia/High-Precision-Event-Synchronization-for-Chronic-Deep-Brain-Stimulation-Recordings), as well as one fully anonymized example LFP and EEG recordings, via the Open Science Framework (OSF) repository (https://osf.io/dnvgb).

**Figure 2.**
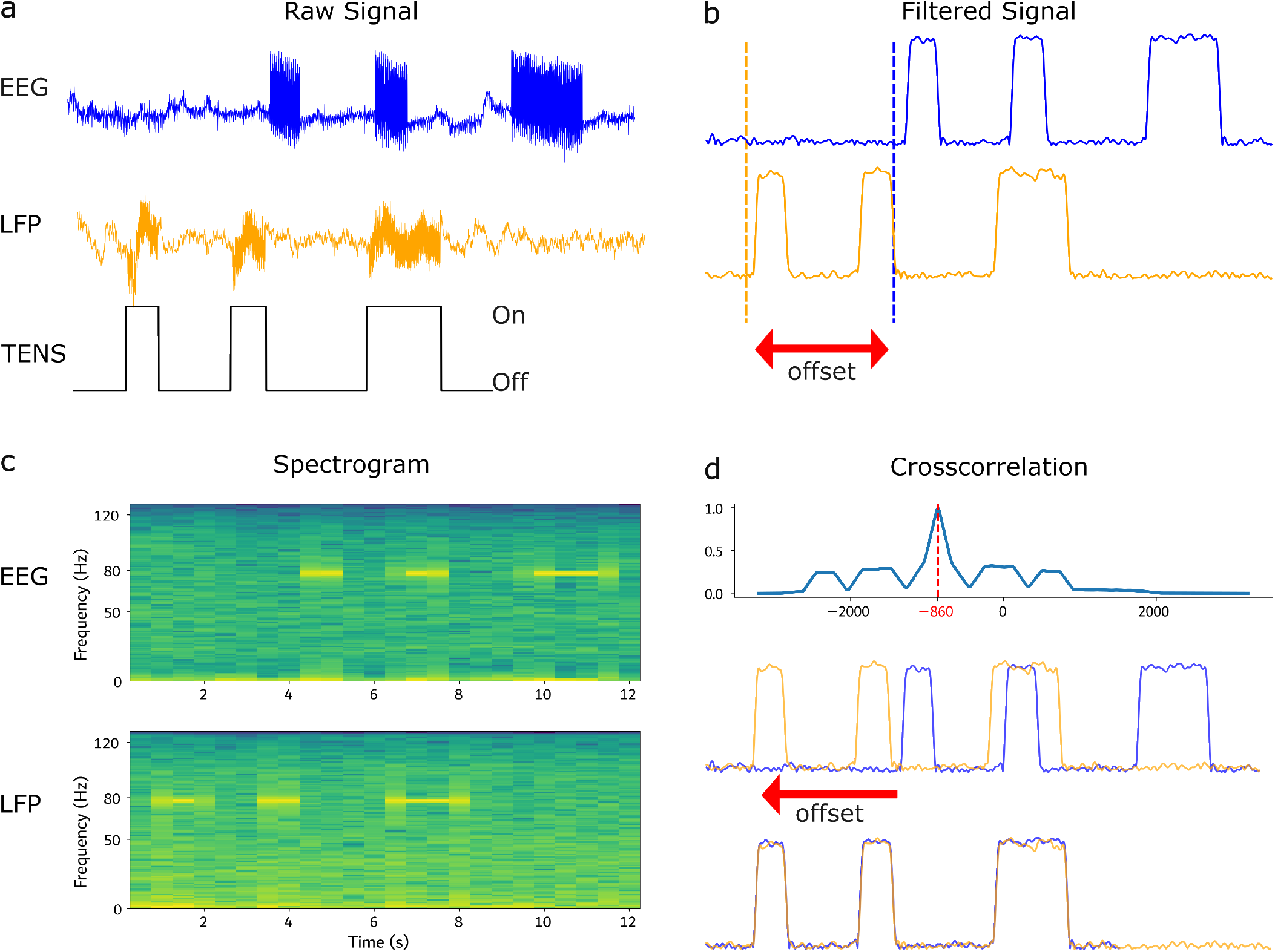
Characterization of induced artifacts and synchronization procedure: a) Raw EEG and LFP data recordings from an example participant are visualized. In this segment, three visible artifacts are induced by repeatedly switching TES stimulation On and Off (as symbolized by the third line). b) Bandpower between 75 and 85 Hz is extracted. The flanks clearly represent the switching of the 80Hz TES stimulation. The visible delay between signals in the EEG and LFP signal represents the offset between the two signals. c) Spectrograms are used to visualize the 80Hz TES artifacts over time. Lighter colors represent a higher power for this time-frequency bin. d) EEG and LFP bandpass-filtered signals are used for alignment. Signals are cross-correlated. The offset with the highest correlation value between the two signals captures the shift between the two signals. By shifting the EEG trace by the computed offset, we can align the two signals, also showing a good visual fit.

### 2.7. Evaluation of synchronization consistency

To assess consistency of our synchronization approach, we used test recordings as well as recordings of a computerized task from our sample participants. For ten participants (P01-P10), recordings to test synchronization were made. First, we wanted to estimate the temporal accuracy of our quantification method in a total of 31 test recordings performed in different recording conditions: *BrainSenseTimeDomain* recording mode with stimulation settings and amplitude used for treatment (abbreviated as **treat**), BrainSenseTimeDomain with stimulation on and amplitude 0 mA (**on**), *BrainSenseTimeDomain* with stimulation off (**o**ff) and *IndefiniteStreamingMode* (**indef**, requires stimulation off). Each test recording consisted of at least 9 artifacts induced by switching the TES stimulation on and off. As all switches were performed manually, induced artifacts can be assumed to be temporally independent from each other. Signals were divided into three non-overlapping groups of at least 3 artifacts each. The LFP and EEG 75-85 Hz power signals were cut into different sub-parts each containing at least 3 artifacts. To ensure that the ground-truth LFP-EEG offset of all sub-parts would be identical, we ensured that corresponding LFP and EEG subparts were cut at the same offset from the first data point. All artifact groups were used separately to compute the offset between the LFP and EEG signal in the previously described optimized cross-correlation approach. By comparing the individually computed offsets, we quantified the respective temporal variation in milliseconds for each recording.

Second, we also tested our synchronization approach for the synchronization of a cognitive Go/NoGo task. During each trial of the task, a fixation cross followed by the letter ‘a’ or ‘A’ was displayed. The more frequent ‘a’ stimulus was the Go cue. Participants were instructed to press the space bar as quickly as possible upon seeing this cue. The stimulus ‘A’ occurred in 30% of trials and represented the NoGo cue. In case of ‘A’, participants were instructed to not press any button. After the space bar was pressed or a timeout of two seconds was reached, the fixation cross was displayed again representing the beginning of the next trial (see Figure 4a). Ten of the participants completed at least one trial of the task (P01-P04 and P06-P11). For synchronization, we induced three artifacts each at the beginning and the end of the recording. To establish the precision of the achieved synchronization, we compare synchronization performed only based on a 50 ms signal sub-part including three TES artifacts at the beginning of the recording versus a 50 ms signal sub-part containing another artifact group at the end of the recording.

**Figure 3.**
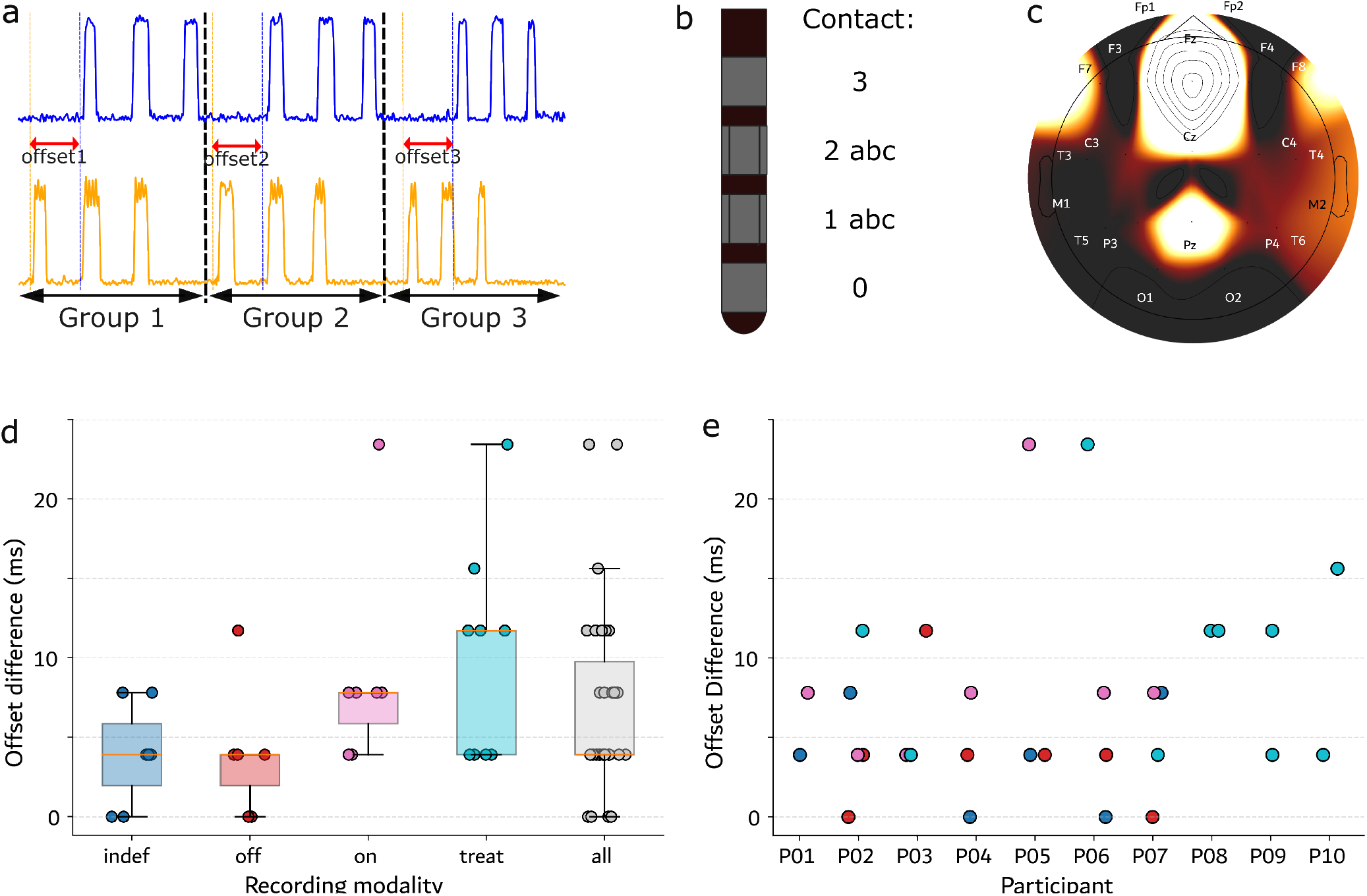
Setup and evaluation of synchronization consistency in test-recordings in 10 participants: a) Artifact test recordings are created by switching the TES stimulator on and off at least 9 times. The test recordings are cut into 3 groups each consisting of at least three artifacts. To evaluate the consistency of computed synchronization offsets for each artifact group, an independent offset (offset1-3) per group is computed and the offsets are compared to determine consistency. b) Electrode layout is visualized. Each electrode tip contains four rings. In the Sensight directional lead, contact rings 1 and 2 are subdivided into 3 different sub-parts. d) Differences between offsets are plotted per recording mode and condition (indef=*IndefiniteStreaming* mode, off=*BrainSenseTimeDomain* mode-stimulation off, on=*BrainSenseTimeDomain*- stimulation on (0 mA), treat= *BrainSenseTimeDomain* mode - stimulation on (stimulation amplitude selected for clinical treatment), all= all recording modes and conditions combined. e) Differences between offsets are plotted per participant. Colors represent recordings mode as labeled in d, c) Channel differences in offset consistency visualized per EEG channel used for synchronization. Darker colors represent lower offset differences across all participants and recording modes. The lowest offset difference is achieved for EEG channels T3,T5 and O2 (8.19 ms) and the highest difference for Fz (49.77 ms).

**Figure 4.**
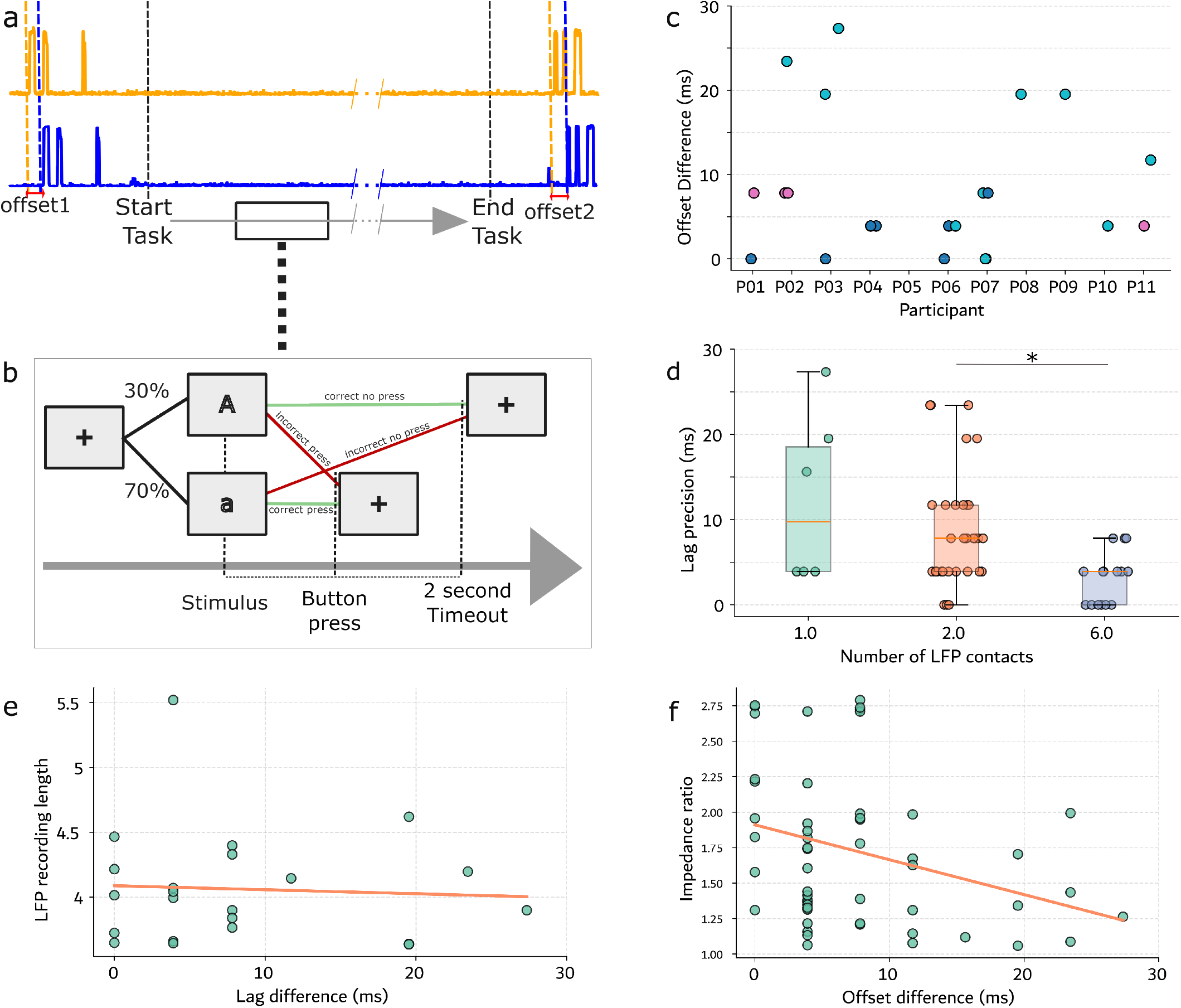
Evaluation of synchronization consistency during a cognitive task. a) At the beginning and the end of the recording, 3 artifacts are induced by switching the TES stimulation on and off. Synchronization, based on the artifact group at the beginning (offset1) and the artifact group at the end (offset2), were compared to calculate synchronization consistency. Artifacts were induced before the task started and after the task ended. b) Task design of the Go/NoGo Task. ‘a’ (70% of cases) and ‘A’ (30% of cases) indicated Go/NoGo Trials, respectively. c) Offset differences in milliseconds per participant. The GoNoGo task was recorded up to four times per participant. Colors represent recording modes (indefinite streaming (dark blue), stimulation on at clinical treatment settings (light blue), stimulation on at 0mA (pink)). d) Offset differences across GoNoGo and synchronization test recordings per number of recording contact pairs. Significant differences are represented by a ‘*’. e) Correlation between recording length of the GoNoGo task and offset difference. f) Correlation between offset difference and maximum impedance per selected contact pair for recordings of the GoNoGo task and synchronization test recordings combined.

### 2.8. Evaluation of additional factors for synchronization consistency

Consecutively, we further evaluated other factors that could influence synchronization consistency. Firstly, using the artifact test recordings, we compared the selection of each EEG channel separately instead of choosing the optimal one out of all EEG channels. This is especially relevant in task settings in which, e.g. due to a recently performed surgery or due to time constraints, only a reduced number of EEG channels can be used. We compute offset consistencies for synchronizing the recording only based on one EEG channel location. Secondly, using the task-data set, we tested for the effect of recording length on synchronization consistency. Prior work has highlighted that an increased recording length (especially above 10 minutes) can result in an increased chance for package loss that could, in turn, effect synchronization consistency [10]. We used Pearson correlation to test for a link between increased recording length and higher offset differences in recordings. For the evaluation of additional factors, we combined recording parameters from synchronization tests and the GoNoGo task. We evaluated a potential connection between recording-contact impedances and synchronization consistency. Based on the circuit configuration, an impedance imbalance between the contacts of the recording pair can increase the contribution of noise that is common to both contacts into the measured signal [21]. Therefore, we hypothesized that a higher impedance imbalance could lead to a stronger TES artifact and, consequently, a better synchronization with less offset inconsistency. To test this hypothesis, we computed the maximal impedance ratio between the two contacts of the optimal recording pair selected for synchronization. To evaluate the impedance *Z* of a Sensight lead ring pair where one ring is divided into sub-parts, we combine the impedance measures of the sub-parts (*Z*1,*Z*2,*Z*3) assuming a parallel setup(1*/Z* = 1*/Z*1 + 1*/Z*2 + 1*/Z*3)) e.g. for contact pair 0-2 (1*/Z*(0 ™2) = 1*/Z*(0™ 2*a*) + 1*/Z*(0™ 2*b*) + 1*/Z*(0™ 2*c*))). The combined impdance is than compared to the impedance of the second recording contact. Lastly, we evaluated whether there was a difference in synchronization consistency dependent on the amount of LFP contact pairs recorded from. We used a linear mixed effects model to test for differences between LFP recording channel numbers in relation to offset consistency while controlling for differences between participants. Post-hoc t-tests allowed for comparing individual electrode number levels.

## 3. Results

### 3.1. Transcutaneous electrical stimulation induced artifacts

Significant differences in 75-85 Hz power between segments including TES artifacts and segments including no artifacts were observed for all LFP and EEG channels. T-values are visualized in 1 b.

### 3.2. Evaluation of alignment consistency for repeated artifacts

Repeated manual on and off switching of TES created multiple consecutive independent artifacts. To assess the reliability of the synchronization method, we tested for consistency of computed offsets between LFP and EEG recordings of three distinct artifact groups, each consisting of at least three TES artifacts. Mean offset difference across all recordings was 7.06 (std=5.92). Offset differences were below 24 ms for all artifact tests in all participants. Notably, during *IndefiniteStreaming* offset differences were below 8 ms with a mean of 3.91 ms (std=3.19) compared to 3.90 ms (std=3.90) during *Brainsense-TimeDomain* stimulation off, 8.93 ms (std=6.66) during *Brain-senseTimeDomain* stimulation on and amplitude set to 0 mA and 10.16 ms (std=6.43) during *BrainSenseTimeDomain* stimulation on at treatment amplitude. Alignment consistency was comparable when instead of optimizing to the best EEG channel, channel O2, T3 or T5 was selected (mean offset=8.19). Only for midline channels Fz (49.77 ms),Cz (10.71 ms) and Pz (12.60 ms), higher offset shifts above 10 ms were observed (compare figure 2 b).

We further evaluated alignment consistency during a cognitive task in 10 participants. Task execution for one run took between 3 and 6 minutes. We show that artifacts induced at the beginning and end of the task, when evaluated separately, provide an alignment with a maximum offset difference of 28 ms (mean=8.35 ms, std=8.29).

### 3.3. Offset-consistency in relation to LFP and EEG channels, recording length and impedance

Further statistical evaluation showed that a higher number of available LFP channels recording synchronization artifacts lead to improved offset-consistency (compare Figure 4 d). A linear mixed-effects model, with participant number as a random effect, revealed a significant effect of number of LFP recording channel pairs on offset difference (*β* = −1.55, *S E* = 0.54, *z* = −2.86, *p* = 0.004, 95% CI interval=[-2.61,-0.49]). Post-hoc t-tests revealed a significant difference between using 2 contact-pairs (as is standard in *BrainSenseTimeDomain* recordings) and using 6 contact pairs (as available in *IndefiniteStream-ing* mode)(*t* = 3.98, *p* = 0.0002). On the other hand, recording length (*mean* = 3.99 minutes, *std* = 0.41, *min* = 3.58, *max* = 5.40) was not significantly correlated to offset differences in our recordings (*r* = −0.06, *p* = 0.80). We found a weak but significant negative correlation between offset differences and impedance ratios of the contacts of the selected LFP contact pair (*r* = −0.31, *p* = 0.02).

## 4. Discussion

### 4.1. TES artifacts as a reliable and precise synchronization method

Overall, TES synchronization proved to be a reliable method for synchronization of EEG and LFP recordings with high consistency. Especially, the high success rate, resulting in significant artifacts and successful synchronization in recordings from all included participants in all EEG and LFP channels stands out (compare Table 3). Therefore, we conclude that this synchronization approach is highly reliable. This method was applicable in 11 out of 14 participant (78.5% success rate). This exceeds reported success rates for methods based on tapping of the IPG, varying between close to 0 [17], 64 [18] or 70% [12] of measurement sessions. Similarly, ECG-based synchronization is also expected to have a lower success rate. To our knowledge, success rate of this method has not systematically been tested. Vivien et al. report visible cardiac artifacts in at least one recording channel in at least one recording in 15 out of 25 participants included in their study [10]. We approximate a success rate of approximately 60% based on this data assuming that their cohort of 25 participants is representative and visible ECG artifacts are a necessary and sufficient condition for successful ECG-based synchronization [10]. At the same time, visibility of ECG artifacts, is dependent on IPG placement [22]. TES based synchronization was a reliable method with high precision in all included participants independent of IPG implantation location. Furthermore, TES synchronization also showed a high synchronization consistency. Both for the artifact test recordings, as well as the cognitive task recordings, our results showed offset differences below 28 ms for all participants, with a mean of below 8.5 ms in both recording types. Benchmark values established for ERP peak-analysis by Williams et al. suggest a cutoff at 16 ms for accurately measuring even the smallest auditory ERP peaks and a cutoff of 46 ms for capturing larger auditory peaks [9]. For 93.5% of our test recordings, and 77.2% of our task recordings, we achieved consistencies below 16 ms while all recordings showed offset differences below 30 ms. These results indicate that TES-based synchronization is suitable for time-domain analysis with the highest precision. TES synchronization also performs well in comparison to other synchronization methods. In the tapping approach, synchronization accuracy is reported to be 570 ms [16]. For stimulation, artifact-based synchronization maximal consistency estimations differ between 8 ms and 16 ms [10, 11]. At the same time, our approach is more versatile across recording modes, enabling recordings during adaptive DBS, where stimulation is adjusted based on LFP signals [23], while manual amplitude switching during recordings is disabled.

**Table 3.** Comparison of synchronization methods. DBS stimulation is abbreviated as stim.

| Synchronization Method | Second recording modality | Mechanism | Reported Success rate | Reported inaccuracy | Possible recording-modes | Notes |
| --- | --- | --- | --- | --- | --- | --- |
| Stimulation artifacts | EEG/MEG | Switching of stim. amplitude | 100% | 8-16 ms | BrainSenseTimeDomain (ON) | automatic cycling possible |
| Tapping on the IPG | video recording | Manual tapping on the IPG | 0-70% | 570 ms | BrainSenseTimeDomain (ON + OFF), IndefiniteStreaming | Success dependent on tapping strength |
| ECG artifacts | ECG | ECG artifacts visible | ≈ 60% | Unknown | BrainSenseTimeDomain (ON) | dependent on IPG implantation location |
| TES artifacts | EEG | low amplitude TES | 78.5% | mean ≤ 8.5 ms | BrainSenseTimeDomain (ON + OFF), IndefiniteStreaming |  |

### 4.2. Particular value of TES synchronization for recordings in IndefiniteStreaming mode

While TES artifact synchronization shows high potential in *BrainSenseTimeDomain* with stimulation ON recordings, the big advantage of TES synchronization lies in its universal usability across recording modes, also when stimulation is switched off. Recordings under stimulation off condition reportedly are less contaminated by ECG artifacts and also do not have any stimulation artifacts [13, 14]. Therefore, these recordings have a higher signal quality and require less artifact removal steps during preprocessing. At the same time, this property makes synchronization using ECG artifacts and stimulation artifacts impossible. On top of that, *IndefiniteStreaming* mode has the unique advantage of allowing for recordings from 3 electrode pairs per implanted side. Our results further suggest, in contrast to the findings by Soh et al. [11], that TES artifact-based synchronization has the highest consistency in *IndefiniteStreaming* mode with all offset differences below 7.8 ms (mean=3.91 ms). This could be due to the low noise levels in recorded LFP signals. Another possible explanation is that the higher number of recording channels gives an advantage potentially allowing for a better pick of an optimal LFP channel with easily detectable TES artifact.

### 4.3. Participant and safety considerations

Generally, we keep TES stimulation low and titrate the highest not unpleasant amplitude below 2mA with all participants. While all included participants could feel the TES, they reported stimulation at 1.5mA or above to not be unpleasant and described it as a light tingling sensation behind the ears. None of the participants experienced any adverse effects of TES stimulation Three participants (not included in our main analysis) found amplitudes above 1.5mA unpleasant. For two of the three participants a stimulation amplitude at 1.24mA could be established that allowed for the generation of visible artifacts in both EEG and LFP recordings. This indicates that, while not optimal, synchronization with lower TES amplitude could still be possible. It is expected that amplitude values above 2mA or with a higher pulse width value would increase artifact amplitudes, which could lead to a further increase in synchronization consistency. However, this was not explored in our study in order to stay at a low and safe threshold.

### 4.4. Further potential factors affecting synchronization consistency

Previously, recording length has been described to potentially cause package loss that could, in turn, affect synchronization accuracy [10]. We hypothesize that our preprocessing method accounting for package loss might partially remedy smaller data losses during data transfer. Indeed, we can not replicate a relationship between recording length and synchronization accuracy for our recording length of up to 5 minutes. It would, however, be interesting to further investigate such a relationship for recordings exceeding 10 minutes. Furthermore, there was a weak negative correlation between impedance imbalance and synchronization offsets in line with our expectations. However, impedance imbalance only has a small effect on offset consistency and also small ratios close to one still allowed for synchronization with low offsets.

## 5. Summary and conclusions

In conclusion, we recommend TES artifact-based synchronization for aligning LFP data recorded with the Medtronic Percept DBS system to EEG signals and task events with high reliability and consistency. The achieved synchronization consistency is suitable for detecting detailed time-domain features. Furthermore, TES-based synchronization proves to be particularly advantageous for recordings under stimulation off in *IndefiniteStreaming* mode as it provides the possibility to leverage ECG-artifact free recordings from all recording-contact pairs for analysis. With regards to EEG recordings, the results confirm that a small number, or even only one recording channel suffice for synchronization, making the approach usable also for recordings directly after surgery or during protocols in which setup time should be minimized. This work supplies a straightforward, reproducible setup with ready-to-use alignment scripts, enabling broader investigations into deep brain electrophysiology via chronic LFP recordings from DBS electrodes.

## Acknowledgements

We want to thank Mirella Waber and Sabine Verkaart for their help with participant planning and overall coordination. We thank Gaetano Leogrando for his technical input regarding the functioning of the Medtronic Percept.

